# Optimization of *arsB* (Acr3) as a positive selection marker in the cyanobacterium *Synechocystis* sp. PCC 6803

**DOI:** 10.1101/2022.09.23.509062

**Authors:** Irene Tejada, Francisco J. Florencio, Luis López-Maury

## Abstract

The use of cyanobacteria as bio-factories for production of numerous compounds of interest (biofuels, bioplastics …) has attracted lots of attention mainly due to their simple nutritional requirements, coupled with the decrease in atmospheric CO_2_ levels. However, although cyanobacteria are easily genetically manipulated, there are few genetic tools developed and, in some cases, the modifications necessary for metabolic engineering are limited for this reason. We have developed a new positive selection marker based on the arsenic resistance system of the cyanobacterium *Synechocystis* sp. PCC 6803. In this cyanobacterium, resistance to arsenic is mediated by the *arsBHC* operon in which *arsB* encodes an arsenite transporter whose mutation confers hypersensitivity to the presence of both arsenite and arsenate. Using *arsB* mutant strain (SARSB) as recipient we introduced plasmids containing both *arsB* and an antibiotic resistance gene and transformants were selected using either arsenic or the antibiotic with similar efficiency. The plasmids and conditions to use the *arsB* gene as a selectable marker have been optimized. Furthermore, we have generated an integrative vector to delete the whole *arsBHC* operon that allows easy introduction of regulated genes in this locus. Analysis of this strain have shown that the *ΔarsBHC* mutant has a higher sensitivity to arsenite than the SARSB strain, even when they are complemented with an *arsB* copy. These suggest that *arsH, arsC* or both could have an additional role in arsenite resistance.

## Introduction

Arsenic is a ubiquitous toxic metalloid that is released to the environment by natural and mankind sources and is likely the most abundant pollutant on Earth (Zhu et al., 2014). It causes serious health problems in many places around the world where the arsenic content in drinking water is well above the recommended limits (Podgorski and Berg, 2020). Arsenic can be found in two biologically active forms, arsenate [As^V^] and arsenite [As^III^], depending on the redox potential of the environment. Arsenate enters the cells through phosphate transporters, as it is a phosphate analogue, and it can replace phosphate in essential biochemical reactions, such as oxidative phosphorylation or glycolysis (Tawfik and Viola, 2011; Kamerlin et al., 2013; Garbinski et al., 2019). The main route for arsenite to enter the cell are aquaglyceroporins (Wysocki et al., 2001; Liu et al., 2002; Meng et al., 2004; Garbinski et al., 2019). Arsenite is more toxic than arsenate and its main toxicity mechanism is through binding to dithiols, forming arsenothiols. Glutathione (GSH) is the main sulphur compound in most cells and arsenite tightly binds to it, forming As^III^-GSH2, and therefore depleting it. This property leads to formation of reactive oxygen species (ROS), as GSH is the main redox buffer in cells. Arsenite binding to proteins, inhibiting them, can also contribute to generate ROS (Flora, 2011). In addition to being toxic, arsenic can also be used by some microorganisms either as an electron acceptor or donor in anaerobic respiratory chain, to grow chemolithotrophically or for anoxygenic photosynthesis carried out by some photosynthetic bacteria and cyanobacteria (Van Lis et al., 2013). Actually, it has been postulated that arsenic compounds played an important role during early life on Earth before the appearance of molecular oxygen as an electron acceptor and donor before the appearance of molecular oxygen (Zhu et al., 2014).

Because of its ubiquitous distribution arsenic resistance is widespread among living organisms where the first line of resistance is based on reduction of arsenate to arsenite, followed by export of the latter outside the cell or its transport to the vacuole (Kruger et al., 2013; Fekih et al., 2018; Garbinski et al., 2019). Arsenite export was probably the first resistance mechanism to evolve as arsenite was dominant form in the ancient Earth (Chen et al., 2020) and is mediated by two proteins families: ArsB proteins, which are present only in bacteria (Rosen, 1999; Chen et al., 2020), and Acr3 proteins, found in different organisms, including bacteria, fungi and plants (Bobrowicz et al., 1997; Rosen, 1999; Garbinski et al., 2019; Chen et al., 2020). Acr3 proteins are older and were probably the first to evolve (Chen et al., 2020). After the appearance of oxygenic photosynthesis and the Great Oxygenation Event arsenate started to accumulate and arsenate reductase appeared (Zhu et al., 2014; Chen et al., 2020). Three families of non-related arsenate reductases have been described: LWPases family of arsenate reductases (arsC2: prototype *Staphylococcus aureus’* ArsC), glutaredoxin-dependent arsenate reductases (arsC1; prototype *Escherichia. coli’s* ArsC) and ArsC2-like arsenate reductases (acr2; prototype *S. cereviseae’s* Acr2) that use thioredoxin, glutaredoxin or mycoredoxin as electron donors (Messens and Silver, 2006; Chen et al., 2020). The arsC2 proteins were the first one to evolve around GOE, when arsenate started to build up on Earth. An additional detoxification mechanism, that is present from bacteria to animals, is arsenic methylation, which covalently adds methyl groups to arsenic and some of these are volatile species that can leave the cell (Ye et al., 2012; Li et al., 2016). Arsenite methylation seems to be ancient, evolving at the same time of Acr3 arsenite export (Chen et al., 2020)

Additional arsenic resistance mechanisms with varying specificities have been recently described all of which seem to have evolved after the GOE (Zhu et al., 2014; Chen et al., 2020). ArsP is a methyl arsenite permease first identified in *Campylobacter jejuni* and is able to confer resistance to methylarsenite (Chen et al., 2015b; Garbinski et al., 2019). Although related to MFS superfamily of transporters it only contains eight transmembrane segments. Three additional MFS superfamily transporters have been identified as arsenic resistance determinants. ArsK is a broad range arsenic exporter being able to confer resistance to arsenite, arsenate, antimonite and methylarsenite. (Shi et al., 2018; Garbinski et al., 2019). An additional MFS protein is frequently associated with ArsK although its substrate specificity has not been identified (Shi et al., 2018; Garbinski et al., 2019). Finally, ArsJ is a MFS transporter involved in arsenate resistance that is usually associated with a *gap* gene, that codes for a GAPDH isoform. This enzyme is able to generate 1-arseno-3-phosphoglycerate which is the substrate for ArsJ (Chen et al., 2016; Garbinski et al., 2019). Three additional arsenic resistance genes related to methylarsenite metabolism have been recently described *arsH*, *arsI* and *arsS*. *arsH* codes for a FMN-oxidoreductase that was described long ago but with no clear *in vivo* function (Neyt et al., 1997; López-Maury et al., 2003; Páez-Espino et al., 2020). More recently it has been shown to be able to oxidise methylarsenite to methylarsenate which is much less toxic (Chen et al., 2015a). *arsI* codes for an methylarsenite lyase that belongs to nonheme iron-dependent dioxygenase family of proteins and catalyse the demethylation of organic arsenite (Yoshinaga and Rosen, 2014; Pawitwar et al., 2017; Yan et al., 2017). *arsS* codes for the first committed step in arsenosugar synthesis which uses the methylarsenite species generated by ArsM as substrate (Xue et al., 2019).

Cyanobacteria are promising bio-factories, since they combine capture and elimination of CO_2_ with bioproduction, in an environmentally friendly way. They have a greater photosynthetic efficiency than higher plants (3-9% for cyanobacteria compared to a maximum of 3% for higher plants) and a faster growth rate, making them ideal for use in biotechnological processes (Hays and Ducat, 2015; Hagemann and Hess, 2018). One of the most promising aspects of the use of these organisms are their simple nutritional requirements, which are limited to sunlight, CO_2_, H_2_O and some microelements with many strains able to use seawater. Cyanobacteria have the advantage that they are easily genetically manipulated so that strains capable of acquiring (or improving) new capabilities can be generated quickly and efficiently by introducing or modifying metabolic pathways. Cyanobacterial strains are available capable of producing a wide variety of compounds of interest (H2, ethanol, isobutyraldehyde, isobutanol, 1-butanol, isoprene, acetone, ethylene, or alkanes (Hays and Ducat, 2015; Hagemann and Hess, 2018). Nevertheless, genome engineering tools for cyanobacteria are currently behind those of model organisms like *E. coli* or yeast and sometime modifications are limited by the lack of suitable selection markers.

In cyanobacteria, arsenic metabolism and resistance has been best studied in the model cyanobacterium *Synechocystis* sp. PCC 6803 (*Synechocystis* hereafter). The main arsenic resistance mechanism is mediated by an operon of three genes (*arsBHC*) that is regulated by an unlinked *arsR* (López-Maury et al., 2003). The operon includes an Acr3-arsenite transporter gene, *arsB*, *arsH*, which codes for a NADPH-FMN reductase, which has been recently proposed to be an methyl-arsenite oxidase although its *in vivo* role in arsenic resistance is not clear (López-Maury et al., 2003; Hervás et al., 2012; Yang and Rosen, 2016), and an arsenate reductase gene, *arsC*. At least two other resistance determinants have been described in *Synechocystis:* an additional arsenate reductase from the *E. coli* family (arsC1 type; encoded by two nearly identical genes, *arsI1* and *arsI2* (López-Maury et al., 2009)) and an arsenite methylase gene, *arsM* (Yin et al., 2011; Xue et al., 2017b). The additional arsenate reductases are only essential for arsenic resistance in the absence of ArsC, probably due to their low expression level (López-Maury et al., 2009). ArsM from several cyanobacteria is able to methylate arsenite to the volatile trimethylarsine using S-adenosyl methionine and glutathione as methyl donors *in vitro* (Yin et al., 2011; Ye et al., 2012), and the *arsM* gene is present in most cyanobacterial genomes. ArsM has a small role in arsenic resistance *in vivo* in cyanobacteria (Xue et al., 2017b) whereas *E. coli* strains carrying *arsM* genes from different cyanobacteria are more resistant to arsenite (Yin et al., 2011; Xue et al., 2017b, 2017a) and provides the substrate for arsenosugars synthesis (Xue et al., 2017b). These compound synthesis is catalised by an additional protein encoded by a neighbour gene, *arsS*, that is co-transcribed with *arsM* (Xue et al., 2017b, 2019). Mutants in *arsM*, but not in *arsS*, are slightly sensitive to arsenite but not to arsenate (Xue et al., 2017b, 2019).

Bioinformatic analyses have shown that arsenic resistance genes are widespread in most sequenced cyanobacterial genomes (Scanlan et al., 2009; Huertas et al., 2014). Some cyanobacterial strains contains gene clusters associated containing some of the following arsenic resistance genes: a periplasmic phosphate binding protein (PstS), a glyceraldehyde-3-phosphate dehydrogenase (Gap3), a major facilitator superfamily transporter (ArsJ) and a homolog of methyl arsenite oxidases, ArsI (Yan et al., 2015), which are probably organized forming an operon with *arsB* (ACR3) and *arsC* genes (Huertas et al., 2014). In some cases, an ArsR coding genes are also localized together with these genes strongly suggesting that they are co-regulated in response to arsenic. The function of these genes in cyanobacteria (with the exception of *arsI* from *Anabaena* sp. PCC 7120; (Yan et al., 2015, 2017)) in relation to arsenic resistance remains to be determined, but it is possible that they will work like their characterised homologs. Furthermore, *arsK* and *arsP* genes homologs can be also identified, some of which are associated together with other arsenic resistance genes (our unpublished observations).

Phosphate plays a central role in arsenate resistance in cyanobacteria given that it controls how much arsenate can enter the cell. Phosphate transport systems are induced by phosphate deprivation (Pitt et al., 2010; Dyhrman and Haley, 2011), therefore increasing arsenate transport. Furthermore, in *Crocosphaera watsonii* it also induces *arsB* expression (Dyhrman and Haley, 2011), while arsenic represses phosphate transport gene expression in *Synechocystis* (Sánchez-Riego et al., 2014). It also affects the redox state of the arsenic species that is accumulated in the media after it is metabolized by cyanobacteria (Zhang et al., 2014). Arsenate accumulates when media contains phosphate but arsenite accumulates in phosphate-depleted media in *Synechocystis* (Zhang et al., 2014).

## Material and methods

### Strains and growth conditions

All *Synechocystis* strains used in this work were grown photoautotrophically on BG11C or BG11C-PO_4_ (BG11C lacking K_2_HPO_4_) at 30° C under continuous illumination (50 μmol photon m^-2^ s^-1^) in 250 mL Erlenmeyer flask containing 50 mL of media. For plate cultures, media was supplemented with 1% (wt/vol) agar. Kanamycin, nourseothricin and spectinomycin were added to a final concentration of 50 μg mL^-1^, 50 μg mL^-1^ and 5 μg mL^-1^, respectively. Plates were supplemented with the indicated amounts of arsenite (NaAsO_2_, Sigma catalogue #S7400) or arsenate (Na_2_HAsO_4_· 7 H_2_O, Panreac catalogue # 131635). Experiments were performed using cultures from the logarithmic phase (0.6-1 OD_750nm_, 3-5 μg chlorophyll mL^-1^; 1 OD_750nm_ is equivalent to 4 10^7^ cells mL^-1^). *Synechocystis* strains are described in Table 1. All *Synechocystis* strains were checked by PCR analysis for complete segregation of all copies of the chromosomes using the appropriated primers as indicated in the figures.

**Table 1.** *Synechocystis* sp. PCC 6803 strains used in this work.

| Strain | Genotype | Mutated ORF | Antibiotic resistance | Reference |
| --- | --- | --- | --- | --- |
| WT | *Synechocystis* sp.<br>PCC 6803 | - | - | Lab stock |
| WT_arsBv1 | *phaAB::arsBv1:Nat<sup>R</sup>* | *slr1993*;<br>*slr1994* | Nat | This work |
| WT_arsBv2 | *phaAB::arsBv2:Nat<sup>R</sup>* | *slr1993*;<br>*slr1994* | Nat | This work |
| SARSB | *arsB::C.K1* | *slr0944* | Km | (López-Maury et al., 2003) |
| SARSB2.1 | *arsB::C.K1*<br>*phaAB::arsBv1:Nat<sup>R</sup>* | *slr0944*;<br>*slr1993*;<br>*slr1994* | Km/Nat/As | This work |
| SARSB2.2 | *arsB::C.K1*<br>*phaAB::arsBv2:Nat<sup>R</sup>* | *slr0944*;<br>*slr1993*;<br>*slr1994* | Km/Nat/As | This work |
| ΔSARSBHC | Δ*arsBHC::aadA* | *slr0944*;<br>*slr0945*;<br>*slr0946* | Sp | This work |
| ΔSARSBHC2.1 | *arsBHC::aadA*<br>*phaAB::arsBv1:Nat<sup>R</sup>* | *slr0944*;<br>*slr0945*;<br>*slr0946*;<br>*slr1993*;<br>*slr1994* | Sp/Nat/As | This work |
| ΔSARSBHC2.2 | *arsBHC::aadA*<br>*phaAB::arsBv2:Nat<sup>R</sup>* | *slr0944*;<br>*slr0945*;<br>*slr0946*;<br>*slr1993*;<br>*slr1994* | Sp/Nat/As | This work |

*E. coli* DH5α cells were grown in Luria Broth supplemented with 100 μg mL^-1^ ampicillin, and 50 μg mL^-1^ spectinomycin when required. For plate cultures, media was supplemented with 1.5% (wt/vol) agar.

### Plasmids construction

Two versions of *arsB* were PCR amplified using VELOCITY DNA Polymerase (Bioline; catalog #BIO-21098) following manufactures recommendations: arsBv1 using oligonucleotides 345 and 346 (arsBv1; 1357 bp, no promoter) and arsBv2 using oligonucleotides 365 y 346 (1515 bp, including the *arsB* promoter region). PCR products digested with *Eco*RI and *Xho*l and cloned in pPHAB_Nat digested with the same enzymes to generate pPHAB_ARSB1 and pPHAB_ARSB2, respectively. Both *arsB* versions were excised using *Hind*III and cloned in *Hind*III digested pRL139 to generate pRLARSB1 and pRLARSB2 (Figure S1;Supplementary data S1). To construct pΔBHC_Sp fragments corresponding to 5’ and 3’ regions of *arsBHC* operon were amplified using oligonucleotides 347-348 (553 bp) and 349-350 (528 bp) respectively. Fragments were joined by overlapping PCR using 347-350 oligonucleotides. The generated fragment was digested *Bss*HII and ligated to pBSSK+ digested in the same way, generating pΔBHC (Figure S2; Supplementary data S2). A 691 bp EcoRI-HindIII fragment from pPHAB_Nat was cloned into pΔBHC to generate pΔBHC-MCS. Finally, 1767 bp PCR amplified fragment containing the *aadA* gene was cloned in pΔBHC-MCS digested with *Hind*III to generate pΔBHC_Sp (Figure S2). Plasmids were sequenced to verify that no mutations were introduced through PCR amplification. All plasmids and oligonucleotides using in this work are described in Tables 2 and 3

**Table 2.**
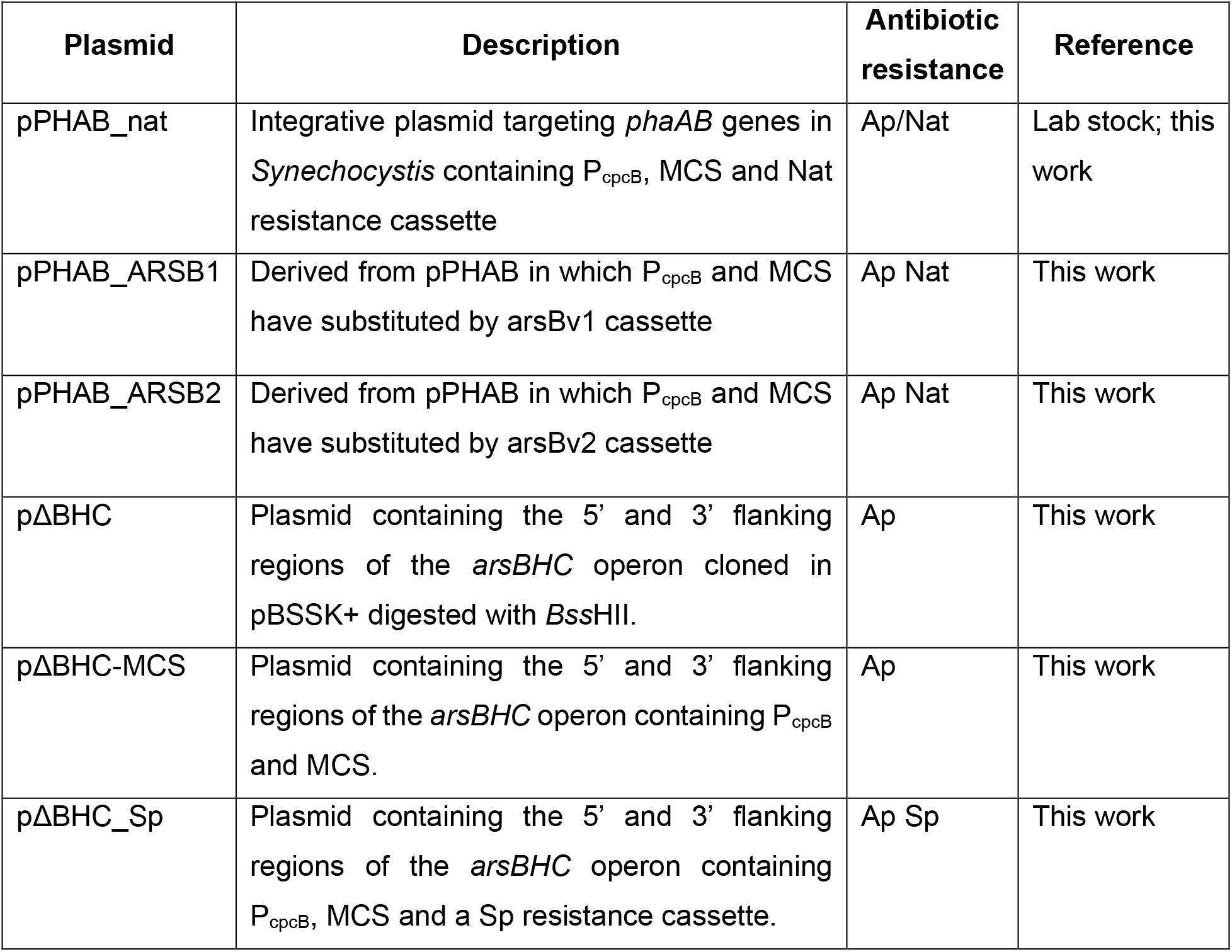
Plasmids used in this work.

**Table 3.**
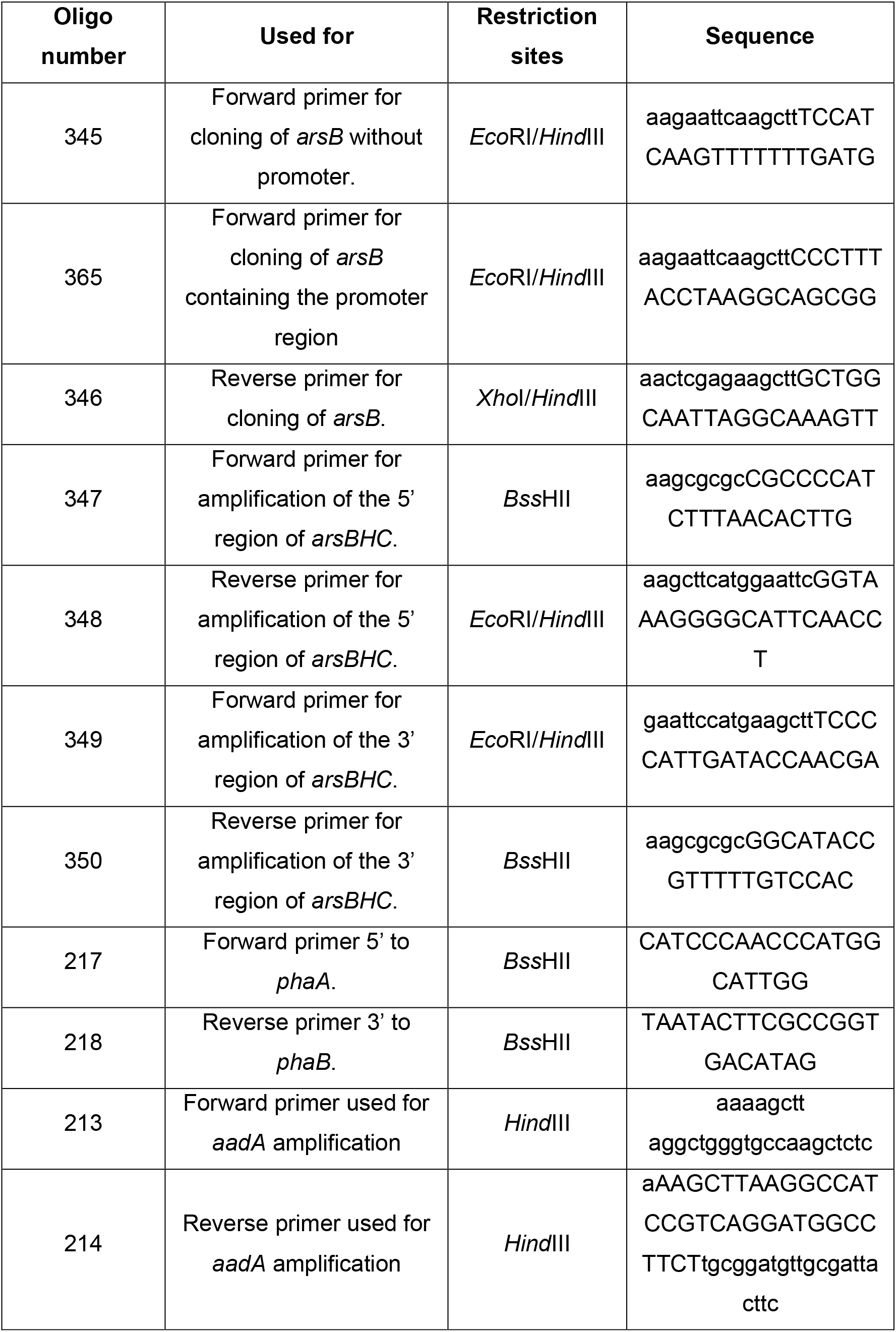
Sequences and uses of the different oligonucleotides used in this work.

### Synechocystis transformation

*Synechocystis* strains were grown in 50 ml of BG11C until exponential phase (OD_750nm_ =0.5-1) and were refreshed daily, at least three time, to keep them between these OD values. Cells were centrifuged at 4500 g for 10 minutes at 25 °C and washed twice with 1 volume of fresh medium. After the second wash, supernatant was discarded, and pellet was resuspended in 1ml of BG11C and adjusted to OD_750nm_= 2.5. 300μL of this cell suspension were transferred to sterile 10 mL transparent polyestyren tubes (Soria Genlab catalog # T1611E), 3 μg of purified DNA were added, and cells were incubated at 30 °C in the light (50 μmol photon m^-2^ s^-1^) for 3 hours with occasional gentle agitation. Cells were plated on Nitrocellulose filters (IMMOBILION-NC, Millipore catalog #HAT08550) placed on BG11C and incubated for 24 h under low light (5 μmol photon m^-2^ s^-1^). Filter were transferred to BG11C plus the selective agent and incubated for 4-6 days in the same conditions until colonies appeared. Colonies were re-streaked at least twice in selective media before PCR analysis for insertion of the constructs and full segregation.

## Results

### Optimization of arsenite concentrations for its use as a selection marker

As a first step we analysed if it was possible to select arsenite resistant cells in an arsenite sensitive background as this is essential to be able to use arsenite as a selective agent. For this WT and SARSB cells were mixed with 1000-fold difference (in the two possible combinations) and serial dilutions were spotted in BG11C, BG11C+Km or BG11C + arsenite. This allowed us to determine if it was possible to select arsenite resistant cells in a mixture of arsenite sensitive and resistant cells. As shown in Figure 1A when WT cells were mixed with a 1000-fold of SARSB cells all dilutions were able to grow in BG11C and BG11C + arsenite but only the first dilutions grew in BG11C+Km as lower concentrations of Km^R^ cells (which correspond to SARSB cells) were present. When the ratio was inverted, all dilutions grew in BG11C+Km and only a few colonies were visible in the first two dilutions in BG11C+arsenite, which represent WT cells (Figure 1A). This showed that was possible to select arsenite resistant colonies in the SARSB background. In order to optimise the use of arsenite for selection we also tested if it could kill cells under the conditions used for *Synechocystis* transformation. This included the use of higher cell concentrations compared to that the one used for the spot assay and nitrocellulose filters to transfer cells form non-selective media to selective media. We carried out the protocol for *Synechocystis’* transformation, in both WT and SARSB strains, and plated the cells directly on plates containing Km or arsenite. We also included one plate in which cells plated on a nitrocellulose filter and were grown for 24 h in BG11C at low light and after 20 h the filter was transferred to an arsenite containing plate. In all cases cells died in the expected media: the WT strain in BG11C+Km and SARSB in BG11C+arsenite although the latter showed some background growth when the nitrocellulose filter was used (Figure 1B).

**Figure 1.**
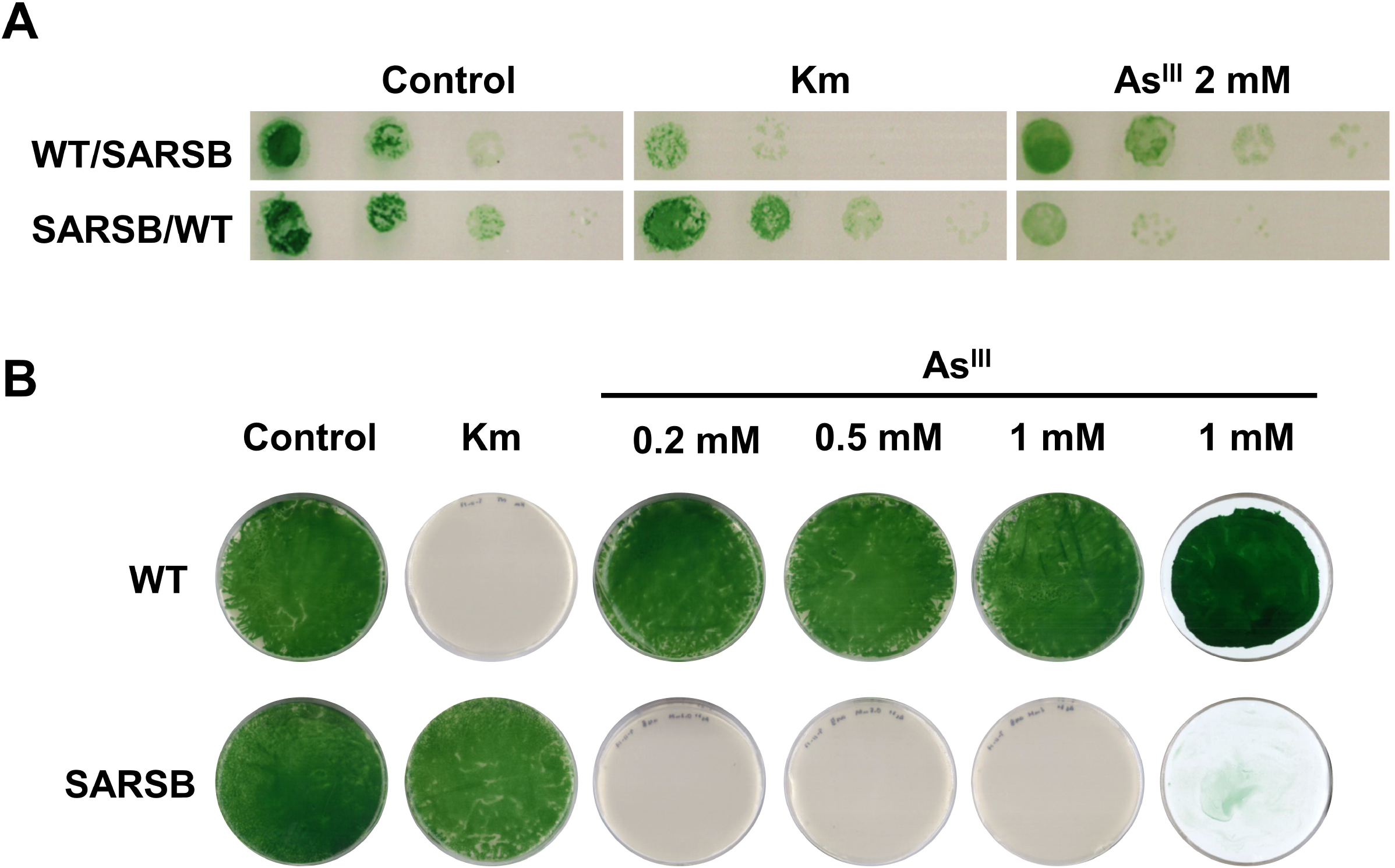
Optimization of arsenite concentration as a selective agent. A. Growth of WT and SARSB mixtures. WT and SARSB from exponentially growing cultures were mixed in 1000:1 (WT/SARSB) and 1:1000 (SARSB/WT) and tenfold serial dilutions of a 1 μg chlorophyll mL^-1^ cells suspension were spotted onto BG11C, BG11C+ Km and BG11C + 1m M arsenite. Plates were photographed after 5 days of growth. B. Growth of WT and SARSB in different arsenite concentrations. 200 μl of exponentially growing cells (OD_750nm_= 0.5 were plated onto BG11C containing Km or the indicated arsenite concentration. Last plate on the right cells were plated on a Nitrocellulose filter that is used in *Synechocystis* transformation protocol.

### Construction of arsenite resistance cassette containing an arsB gene

In order to be able to use arsenite as a selection agent an arsenite resistance gene is needed and for that we have amplified two version of *Synechocystis’ arsB* gene: one without the native promoter (arsBv1) and one including the *arsB* native promoter (arsBv2). Both genes were PCR amplified with oligonucleotide that included *HindIII, Eco*RI and *Xho*I restriction sites that allowed us to clone them in the desired plasmid to test whether they will confer resistance in *Synechocystis*. Both versions were cloned in pPHAB_Nat between *Eco*RI and *XhoI* sites to generate integrative plasmids (targeting phaAB operon involved in polyhydroxybutyrate synthesis) containing both the arsenite (*arsB*) and Nat (nourseothricin) resistance cassettes generating pPHAB_ARSB1 and pPHAB_ARSB2 (Figure 2A). These plasmids integrated into *phaAB* genes in *Synechocystis’* genome. We have also cloned both versions in pRL139 (Elhai and Wolk, 1988) in order to generate a cassette with symmetric MCS at both ends allowing rapid subcloning of it (pRLARSB1 and pRLARSB2; Figure S1; supplementay data S1).

**Figure 2.**
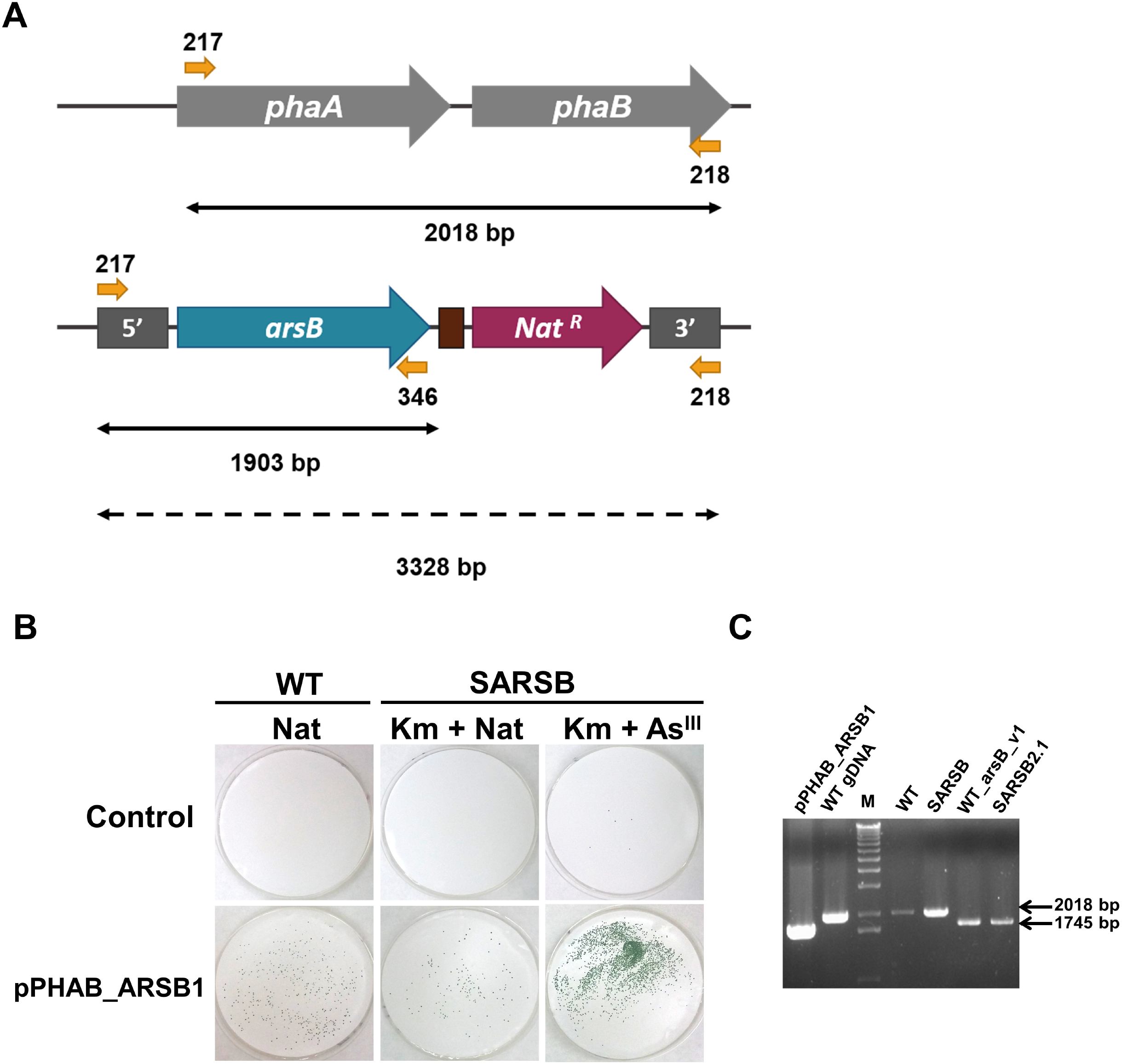
Selection of arsenite resistant colonies in SARSB. A. Schematic representation of the genomic region in which *arsB* and Nat cassettes were introduced in WT and transformed strains. B. Selection of arsenite/Nat resistant colonies in WT and SARSB strains. Both strains were transformed with pPHAB_ARSB1 or with now DNA (control) and plated in BG11C+Nat (WT), BG11C+Km+Nat or BG11C+Km+2 mM arsenite. Plates were photographed after 10 days of growth, C. PCR analysis of the *phaAB* locus in WT, SARSB, WT_arsBv1 and SARSB2.1 strains using oligonucleotides 217, 218 and 346. The band amplified using 217-218 is not amplified in the mutant version because the extension time used was shorter than the required for amplification. WT gDNA and pPHAB_ARSB1 plasmid were included as controls.

### Arsenite selection in a SARSB strain

Both WT and SARSB strains were transformed with both pPHAB_ARSB1 and pPHAB_ARSB2. Transformants were selected using either Nat (WT and SARSB) or 2 mM arsenite (SARSB). Colonies appeared in both strains in Nat containing plates but also in arsenite plates in the case of SARSB (Figure 2B). Correct integration of the plasmids was verified by PCR analysis in both cases (Figure 2C). Colonies segregated independently of in which media they were selected, demonstrating that arsenite can be used for selection in the SARSB background.

To further expand the genetic toolbox in *Synechocystis* a plasmid carrying *arsBHC* flanking regions together with a MCS and spectinomycin/streptomycin resistance cassette was generated (Figure 3 and Figure S2). This will allow introduction of regulated genes in *Synechocystis’* genome in the *arsBHC* locus and later use of the *arsB* as arsenite resistance gene. After transforming *Synechocystis* WT with plasmid pΔBHC_Sp a strain deleting the *arsBHC* operon was generated (ΔSARSBHC; Figure 3). This new strain was transformed with plasmids pPHAB_ARSB1 and pPHAB_ARSB2 and colonies were selected using either Nat or 2 mM arsenite. Colonies appeared in Nat containing plates but not in arsenite containing plates, suggesting that the ΔSARSBHC was more sensitive to arsenite than the SARSB strain. Transformation was repeated with pPHAB_ARSB2 and 1 mM arsenite as selective media. Colonies appeared in both media but with lower frequency in the arsenite plates (Figure 3), strongly suggesting that the ΔSARSBHC is in fact more sensitive to arsenite (see below). In both cases correct integration and full segregation of the mutation was checked by PCR analysis with similar results to SARSB. Further optimization will be needed to determine an optimal concentration of arsenite in the ΔSARSBHC strain.

**Figure 3.**
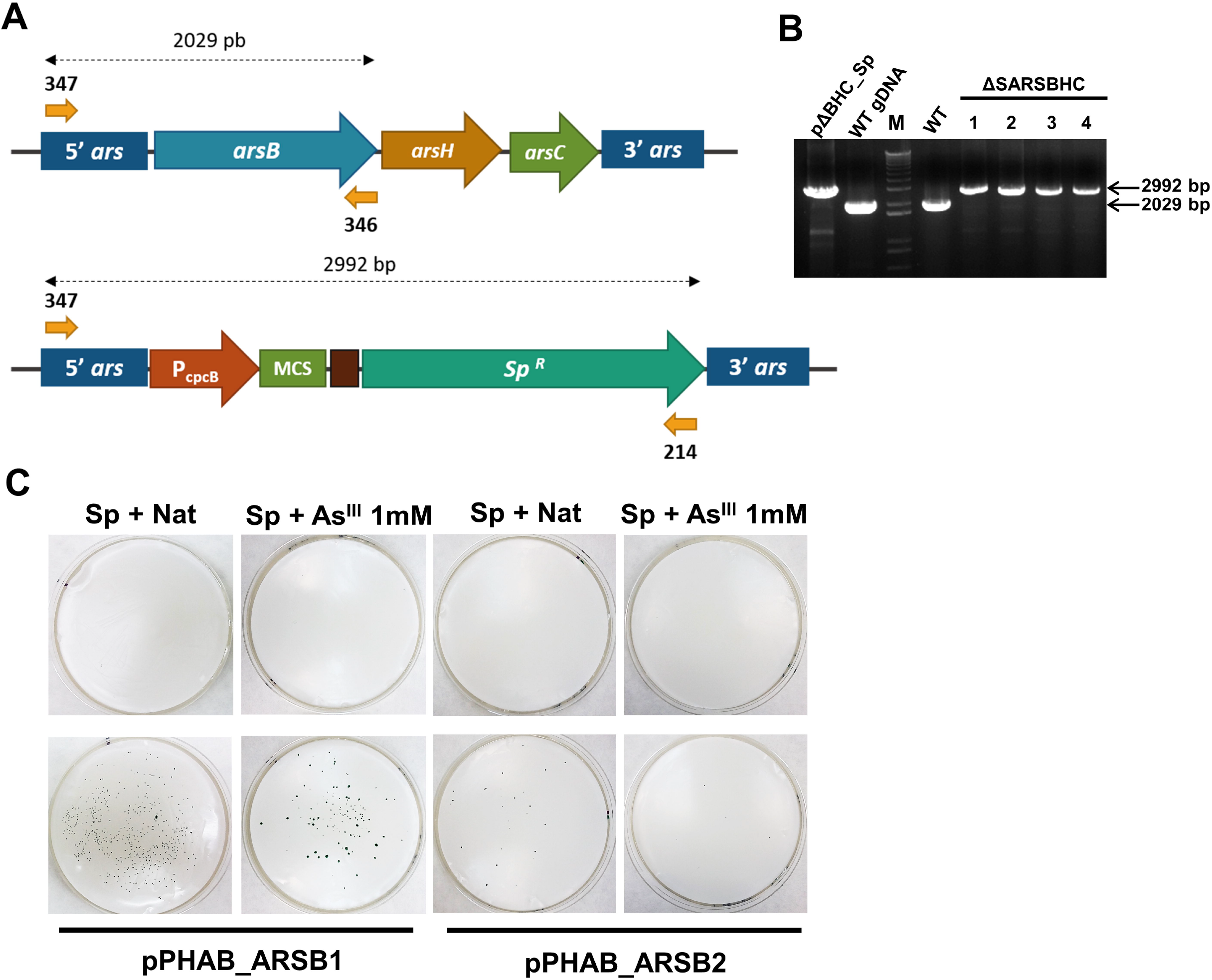
Construction of ΔSARSBHC strain. A. Schematic representation of the *arsBHC* genomic region in WT and ΔSARSBHC strains. Oligonucleotides used for PCR analysis and the expected sizes of the amplified bands are also shown. B. PCR analysis of the *arsBHC* locus in WT and ΔSARSBHC strains using oligonucleotides 347, 346 and 214. WT gDNA and pΔBHC_Sp plasmid were included as controls. C. Selection of arsenite/Nat resistant colonies in the ΔSARSBHC strain. The strain was transformed with pPHAB_ARSB1 or pPHAB_ARSB2 and plated in BG11C+Sp+Nat or BG11C+Sp+1 mM arsenite. Plates were photographed after 10 days of growth.

### ΔSARSBHC strain is extremely sensitive to arsenic

In order to study arsenic resistance in these new strains serial dilutions of the strains were plated in arsenite or arsenate containing plates. As is shown in Figure 4, SARSB and ΔSARSBHC strains were extremely sensitive to both arsenate and arsenite. The ΔSARSBHC was more sensitive to both arsenic forms than SARSB (Figure 4) as it was unable to grow in the presence of 50 μM of arsenite and was much more sensitive to 50 μM and 0.1 mM arsenate, displaying yellowish appearance in these concentrations.

**Figure 4.**
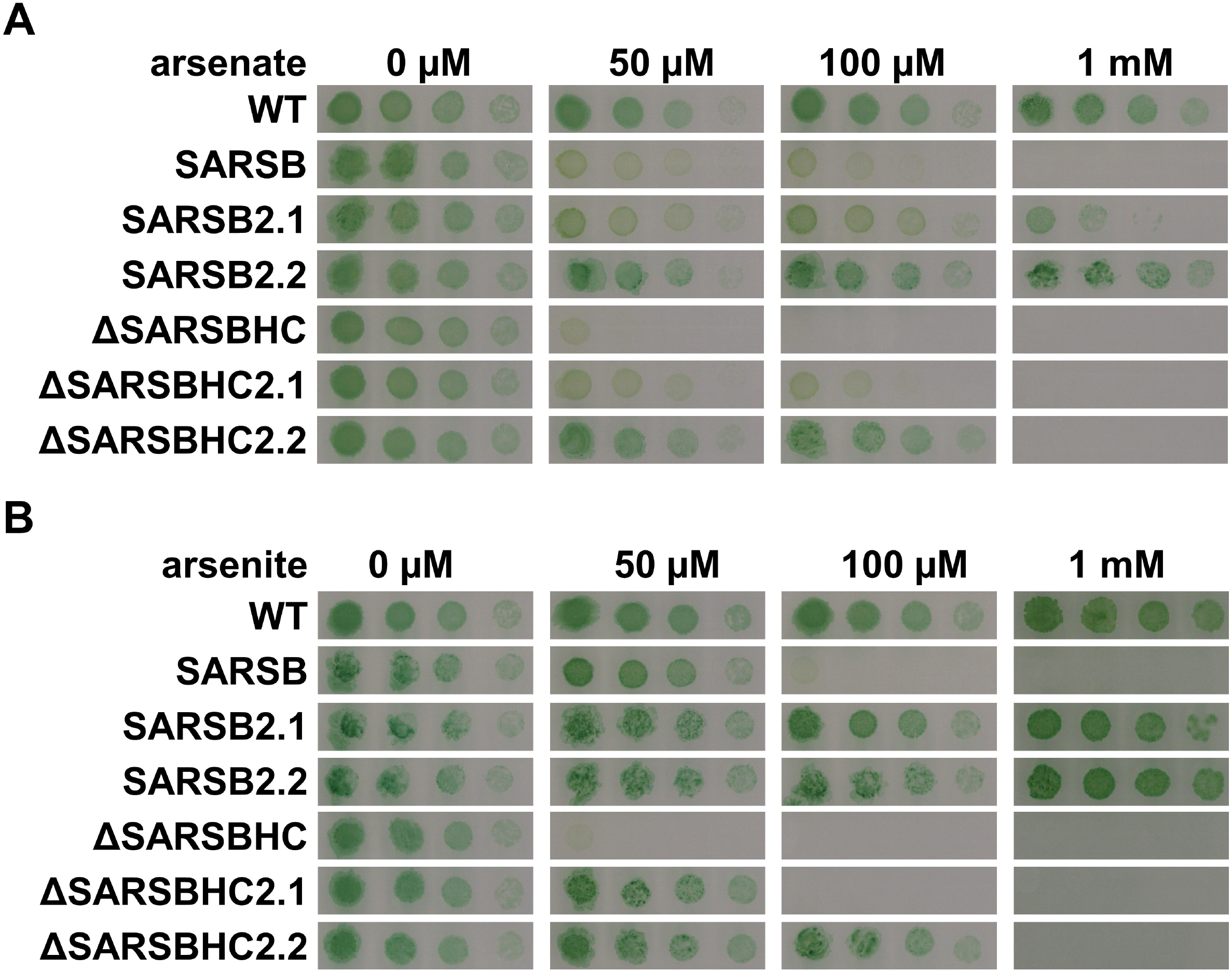
Phenotypic characterisation of ΔSARSBHC and complemented strains. A. Growth of WT, SARSB, SARSB2.1. SARSB2.2, ΔSARSBHC, ΔSARSBHC2.1, ΔSARSBHC2.2 strains in different arsenate concentrations. Tenfold serial dilutions of a 1 μg chlorophyll mL^-1^ cells suspension were spotted onto BG11C-PO4 containing the indicated arsenate concentrations. Plates were photographed after 5 days of growth. B. Growth of WT, SARSB, SARSB2.1. SARSB2.2, ΔSARSBHC, ΔSARSBHC2.1, ΔSARSBHC2.2 strains in different arsenite concentrations. Tenfold serial dilutions of a 1 μg chlorophyll mL^-1^ cells suspension were spotted onto BG11C containing the indicated arsenite concentrations. Plates were photographed after 5 days of growth.

We have also analysed these two strains when they were complemented with both versions of the *arsB* cassette (SARSB2.1, SARSB2.2, ΔSARSBHC2.1 and ΔSARSBHC2.2, respectively) for their resistance to arsenate and arsenite. The complemented strains were more resistant to both arsenate and arsenite then their parental strains (Figure 4). Nevertheless, the levels of resistance varied depending on the background and the version of the *arsB* cassette. ΔSARSBHC derived strains were more sensitive than SARSB strains both to arsenate and arsenite suggesting that either *arsH* or *arsC* might have a role in resistance to both arsenic forms. In addition, in both backgrounds, arsBv1 cassette conferred less resistance to arsenic than arsBv2 cassette. This suggests that higher expression in response to arsenic presence that is predicted for arsBv2 cassette confers and advantage to low and non-inducible expression in arsBv1 cassette.

## Discussion

The use of arsenite as an additional positive selection marker in *Synechocystis* (and probably in other cyanobacteria) will allow to generate strains with up to 6-7 modifications using positive selection with already in use antibiotics (Taton et al., 2014), although it needs strains lacking at least *arsB*. Furthermore, if combined with a strain with *arsB* or *arsBHC* deleted it can be used to generate strains using positive selection without the use of antibiotics and antibiotic resistance genes. If these deleted strains are generated without the use antibiotic resistance, using *sacB* or *mazF* counterselection or CRISPR (Behler et al., 2018), will allow the generation of non-GMO strains that can be very useful for biotechnological uses. Although we have observed a marked difference in the resistance levels for the two versions of the *arsB* cassette constructed (Figure 4) both can be used for positive selection (Figures 2 and 3). This difference could be easily explained by the levels of expression expected in both: low and constitutive in arsBv1 and inducible and transient strong expression for arsBv2. Finally, it is also possible that induction of genes under P_arsB_ promoter in these strains is much more sensitive to the presence of arsenic, allowing the use of very low concentrations, below the permitted levels, to induce the genes under its control in large scale production systems making these systems more flexible.

The ΔSARSBHC strain is more sensitive to arsenic than the SARSB strain and although the sensitivity to arsenate could be easily explained by the lack of both *arsC* (the main arsenate reductase) and *arsB* (López-Maury et al., 2003, 2009), but the higher sensitivity to arsenite of ΔSARSBHC2.1 and ΔSARSBHC2.2 points to an additional role of either *arsC, arsH* or both in arsenite resistance. Given that ArsH has been described as methylarsenite oxidase (Chen et al., 2015a), it is possible that this sensitivity is due to lower oxidation of the methylarsenite species generated by ArsM in a strain lacking also ArsH. Methylarsenite species are extremely toxic compounds that have been postulated to be ancients forms of antibiotics (Li et al., 2016; Chen et al., 2019) and they may accumulate to high concentration in this background. ArsH has also been proposed to play a role in protection against oxidative stress generated by arsenite in *Synechocystis* and *Pseudomonas putida* (Hervás et al., 2012; Páez-Espino et al., 2020) and therefore it is possible that the lack of ArsH may cause increased ROS accumulation in these strains. Another possibility is that the futile cycle of arsenite oxidation/reduction, that normally occurs in *Synechocystis*, is generating more arsenate that can not be efficiently detoxified in this strain due to the lack of *arsC* (Zhang et al., 2014). However, this is unlikely as there are 2 additional genes coding for an arsenate reductase in *Synechocystis* genome and the concentration of arsenate used is lower compared to the ones inhibiting growth of *arsC*^-^ strains or the strains lacking all three genes for arsenate reductases (López-Maury et al., 2003, 2009). The ΔSARSBHC strain will be also valuable to explore the function of *arsMS* in arsenite resistance as these have only moderate effect in resistance in a WT background (Xue et al., 2017b, 2019), probably because of the presence of *arsBHC*. It would be interesting to analyse if expression of *arsMS*, alone or together with *arsH*, under the control of an arsenic inducible promoter, such as P_arsB_, can confer higher levels of arsenic resistance.

Finally, the ΔSARSBHC strain will very useful to explore/test the function of the new putative arsenic resistance genes found in cyanobacteria, which have been poorly characterised *in vivo* (Huertas et al., 2014). Even though some of these genes have been used to complement hypersensitive *E. coli* strains to characterise their function (Yin et al., 2011; Xue et al., 2017b, 2019; Yan et al., 2017), cyanobacteria possess completely different metabolism and cell organization. For example, when analysing membrane transporters or redox proteins the results might be not clear in *E. coli* due the lack of proper targeting to the membranes (thylakoid or plasma membrane in cyanobacteria; (Mullineaux and Liu, 2020)) or the lack of correct redox partners which markedly differs between *E. coli* and cyanobacteria (Florencio et al., 2006).

## Acknowledgments

This research was funded by grant numbers BIO2016-75634-P and PID2019-104513GB-I00 from the Ministerio de Economía y Competividad (MINECO) and the Agencia Estatal de Investigación (AEI), respectively, and by Junta de Andalucía Group BIO-284, co-financed by European Regional Funds (FEDER), to Francisco J. Florencio. I.T. was recipient of a Beca de Iniciación a la Investigación from VI plan propio de investigación y transferencia Universidad de Sevilla

