## Supplementary data S1 for "Optimization of *arsB* (Acr3) as a positive selection marker in the cyanobacterium *Synechocystis* sp. PCC 6803"

Sequences of inserts cloned in pRLARSB plasmids

>pRLARSBv1

gaattcgagctcggtacccggggatcctctagagtcgacctgcaggcatgcaagcttTCCATCAAGTTTTTTTGATGAGTTGTCATGGTCAATCGCATTAACCCCAAAGCCATTAAGGCTGGGGGAACGCTCAATTTGTTTGAAAAATACCTTACCCTCTGGGTTGCCCTTTGTATCGTCATTGGCATTGCCCTGGGGAAGTTATTGCCCGCGGTGGCCCAAACCCTCGATTCCTGGAGCATTTATAACGTTTCCATTCCGATCGCCATTTGCTTGTTTTTCATGATGTACCCCATCATGGTGAAAATTGATTTTTCCCAGGCACGACAAGCGGTCAAGGCTCCTAAGCCGGTGATATTGACCCTGGTGGTGAATTGGGTAATTAAACCGTTTACGATGGTGATTTTTGCCCAGTTTTTCCTGGGTTATCTTTTCGCCCCGTTGTTGACTGCGACGGAGATAATCCGGGGGCAAGAGGTAACCTTAGCCAATTCCTACATTGCCGGTTGCATTTTGCTCGGTATTGCGCCCTGTACCGCCATGGTATTGATGTGGGGTTACCTTTCCTATAGCAATCAGGGTTTAACCTTGGTGATGGTGGCGGTCAATTCTTTGGCTATGCTATTCCTTTACGCACCTTTGGGTAAATGGCTGTTGGCGGCCAGTAACTTGACAGTGCCTTGGCAAACCATTGTCTTATCGGTGTTAATTTACGTGGGTTTGCCCCTGGCCGCCGGTATTTATAGTCGTTACTGGATTTTGAAACATAAGGGACGGCAATGGTTTGAGAGTCAATTTTTGCATTACCTCAGCCCGATCGCCATTGTGGCTTTGTTATTGACGTTGATTTTGCTCTTTGCTTTTAAGGGGGAATTGATTGTTAATAATCCTCTGCATATTTTTCTAATTGCCGTGCCGTTATTTATCCAAACCAATTTCATCTTTCTCATTACCTATGTTCTGGGGTTAAAACTCAAACTCAGTTACGAAGATGCCGCCCCCGCCGCCCTGATCGGTGCCAGTAATCATTTTGAAGTGGCGATCGCCACCGCTGTGATGTTATTTGGCCTCAATTCTGGAGCTGCTTTGGCCACCGTAGTGGGGGTATTAATTGAAGTGCCAGTGATGTTAATGCTGGTGGAAATTTGCAAAAAAACAGCCTTTTGGTTCCCCAGGGACCCGGAAAAAGCCACTTTGCTAGACCCCCGTTGTATAAATCAAGAAATTAGGATCTAAAAACGTGTTTGGAAAGTTTTATTCTGCCCCCTCAATCGACCAACTAAAGCGACCGCAATACTGGGGAACTTTCACTCAGTTTCCCCCCAAATTTGGGGGGCCAGGGGGGCTTTTAAAACAAGCTCTAACCAACTTTGCCTAATTGCCAGCaagcttgcatgcctgcaggtcgactctagaggatccccgggtaccgagctcgaattc

>pRLARSBv2

gaattcgagctcggtacccggggatcctctagagtcgacctgcaggcatgcaagcttCCCTTTACCTAAGGCAGCGGAGGATGTGAATGGAACAATGGGGGCAATGAGATTTCCATAAAGTACTCTATTTTGCTCCTTTTCCATCGGAAGACACGGCAACAGTAGACCGGGGACAGACCGAGTTGACATATCAAGTTTTTTTGATGTAAGATTATTCCATCAAGTTTTTTTGATGAGTTGTCATGGTCAATCGCATTAACCCCAAAGCCATTAAGGCTGGGGGAACGCTCAATTTGTTTGAAAAATACCTTACCCTCTGGGTTGCCCTTTGTATCGTCATTGGCATTGCCCTGGGGAAGTTATTGCCCGCGGTGGCCCAAACCCTCGATTCCTGGAGCATTTATAACGTTTCCATTCCGATCGCCATTTGCTTGTTTTTCATGATGTACCCCATCATGGTGAAAATTGATTTTTCCCAGGCACGACAAGCGGTCAAGGCTCCTAAGCCGGTGATATTGACCCTGGTGGTGAATTGGGTAATTAAACCGTTTACGATGGTGATTTTTGCCCAGTTTTTCCTGGGTTATCTTTTCGCCCCGTTGTTGACTGCGACGGAGATAATCCGGGGGCAAGAGGTAACCTTAGCCAATTCCTACATTGCCGGTTGCATTTTGCTCGGTATTGCGCCCTGTACCGCCATGGTATTGATGTGGGGTTACCTTTCCTATAGCAATCAGGGTTTAACCTTGGTGATGGTGGCGGTCAATTCTTTGGCTATGCTATTCCTTTACGCACCTTTGGGTAAATGGCTGTTGGCGGCCAGTAACTTGACAGTGCCTTGGCAAACCATTGTCTTATCGGTGTTAATTTACGTGGGTTTGCCCCTGGCCGCCGGTATTTATAGTCGTTACTGGATTTTGAAACATAAGGGACGGCAATGGTTTGAGAGTCAATTTTTGCATTACCTCAGCCCGATCGCCATTGTGGCTTTGTTATTGACGTTGATTTTGCTCTTTGCTTTTAAGGGGGAATTGATTGTTAATAATCCTCTGCATATTTTTCTAATTGCCGTGCCGTTATTTATCCAAACCAATTTCATCTTTCTCATTACCTATGTTCTGGGGTTAAAACTCAAACTCAGTTACGAAGATGCCGCCCCCGCCGCCCTGATCGGTGCCAGTAATCATTTTGAAGTGGCGATCGCCACCGCTGTGATGTTATTTGGCCTCAATTCTGGAGCTGCTTTGGCCACCGTAGTGGGGGTATTAATTGAAGTGCCAGTGATGTTAATGCTGGTGGAAATTTGCAAAAAAACAGCCTTTTGGTTCCCCAGGGACCCGGAAAAAGCCACTTTGCTAGACCCCCGTTGTATAAATCAAGAAATTAGGATCTAAAAACGTGTTTGGAAAGTTTTATTCTGCCCCCTCAATCGACCAACTAAAGCGACCGCAATACTGGGGAACTTTCACTCAGTTTCCCCCCAAATTTGGGGGGCCAGGGGGGCTTTTAAAACAAGCTCTAACCAACTTTGCCTAATTGCCAGCaagcttgcatgcctgcaggtcgactctagaggatccccgggtaccgagctcgaatt

c
