## Supplementary data S2 for "Optimization of *arsB* (Acr3) as a positive selection marker in the cyanobacterium *Synechocystis* sp. PCC 6803"

Sequence of pΔBHC_Sp

>pΔBHC_Sp

aagcgcgcCGCCCCATCTTTAACACTTGTCAGTAATTGCTGAAATTGTTTGACTCCAGCGCCGGTAATACCAAGGTTGTATAGACTGATCTCTGGGCGATCCAAACGTCGCCTTAGTTGACTGATCCAGGACTGATTGCCCCCATCTCCGGCGGTGAAAGAGTCACCGACAAACACAACTTTTTTGCCGGGGGCGGAATCAAAATTGTAGTCTACGTCATCTACAAAACCGAGGTTATTGGTGCCATAGACAATGTCGTAATCAACTTCATCCCCATAGATAGCAACACTTCTAATTTTCGTTCTTGGCTCATACCTAACAGTTTTATCCCCCGTTCTCACCAGGGAAGGCTTGGTAAAAAATAAAATTCGGTCATAGAGACTGTATTTACCTATGCCAGTTAACCTCAATGCAATTTCAGCTATTACCAAACTAAGGACAATTGAAGCAAGAATTAAAGCAGTATTTTGACCAATTTTAATAATTTTGGGGGCGGGCATTGTTTTATCAGAAAAATAGGTTGAATGCCCCTTTACCgaattcCACCTGTAGAGAAGAGTCCCTGAATATCAAAATGGTGGGATAAAAAGCTCAAAAAGGAAAGTAGGCTGTGGTTCCCTAGGCAACAGTCTTCCCTACCCCACTGGAAACTAAAAAAACGAGAAAAGTTCGCACCGAACATCAATTGCATAATTTTAGCCCTAAAACATAAGCTGAACGAAACTGGTTGTCTTCCCTTCCCAATCCAGGACAATCTGAGAATCCCCTGCAACATTACTTAACAAAAAAGCAGGAATAAAATTAACAAGATGTAACAGACATAAGTCCCATCACCGTTGTATAAAGTTAACTGTGGGATTGCAAAAGCATTCAAGCCTAGGCGCTGAGCTGTTTGAGCATCCCGGTGGCCCTTGTCGCTGCCTCCGTGTTTCTCCCTGGATTTATTTAGGTAATATCTCTCATAAATCCCCGGGTAGTTAACGAAAGTTAATGGAGATCAGTAACAATAACTCTAGGGTCATTACTTTGGACTCCCTCAGTTTATCCGGGGGAATTGTGTTTAAGAAAATCCCAACTCATAAAGTCAAGTAGGAGATTAATTCAgtcgactctagacatatgggatccgcggccgcgatatcctcgagCCAGGCATCAAATAAAACGAAAGGCTCAGTCGAAAGACTGGGCCTTTCGTTTTATCTGTTGTTTGTCGGTGAACGCTCTCaaagcttaggctgggtgccaagctctcgggtaacatcaaggcccgatccttggagcccttgccctcccgcacgatgatcgtgccgtgatcgaaatccagatccttgacccgcagttgcaaaccctcactgatccgcatgcccgttccatacagaagctgggcgaacaaacgatgctcgccttccagaaaaccgaggatgcgaaccacttcatccggggtcagcaccaccggcaagcgccgcgacggccgaggtcttccgatctcctgaagccagggcagatccgtgcacagcaccttgccgtagaagaacagcaaggccgccaatgcctgacgatgcgtggagaccgaaaccttgcgctcgttcgccagccaggacagaaatgcctcgacttcgctgctgcccaaggttgccgggtgacgcacaccgtggaaacggatgaaggcacgaacccagtggacataagcctgttcggttcgtaagctgtaatgcaagtagcgtatgcgctcacgcaactggtccagaaccttgaccgaacgcagcggtggtaacggcgcagtggcggttttcatggcttgttatgactgtttttttggggtacagtctatgcctcgggcatccaagcagcaagcgcgttacgccgtgggtcgatgtttgatgttatggagcagcaacgatgttacgcagcagggcagtcgccctaaaacaaagttaaacatcatgagggaagcggtgatcgccgaagtatcgactcaactatcagaggtagttggcgtcatcgagcgccatctcgaaccgacgttgctggccgtacatttgtacggctccgcagtggatggcggcctgaagccacacagtgatattgatttgctggttacggtgaccgtaaggcttgatgaaacaacgcggcgagctttgatcaacgaccttttggaaacttcggcttcccctggagagagcgagattctccgcgctgtagaagtcaccattgttgtgcacgacgacatcattccgtggcgttatccagctaagcgcgaactgcaatttggagaatggcagcgcaatgacattcttgcaggtatcttcgagccagccacgatcgacattgatctggctatcttgctgacaaaagcaagagaacatagcgttgccttggtaggtccagcggcggaggaactctttgatccggttcctgaacaggatctatttgaggcgctaaatgaaaccttaacgctatggaactcgccgcccgactgggctggcgatgagcgaaatgtagtgcttacgttgtcccgcatttggtacagcgcagtaaccggcaaaatcgcgccgaaggatgtcgctgccgactgggcaatggagcgcctgccggcccagtatcagcccgtcatacttgaagctagacaggcttatcttggacaagaagaagatcgcttggcctcgcgcgcagatcagttggaagaatttgtccactacgtgaaaggcgagatcaccaaggtagtcggcaaataatgtctaacaattcgttcaagccgacgccgcttcgcggcgcggcttaactcaagcgttagatgcactaagcacataattgctcacagccaaactatcaggtcaagtctgcttttattatttttaagcgtgcataataagccctacacaaattgggagatatatcatgaaaggctggctttttcttgttatcgcaatagttggcgaagtaatcgcaacatccgcaAGAAGGCCATCCTGACGGATGGCCTTAAGCTTTCCCCATTGATACCAACGATACCAAACCCCCAATTTGGCTATATGCTTAGGGAAATGGCTTTTTGCCACGAAACTTCACTAATCGTTATACAAGTTAATCAAAGGAGGGTTGTGGATCTTGGAAACCAATAAAGAAAGGATATTGGTGGTCGATGACGAGGCCAGCATCCGACGCATACTGGAAACCCGTTTATCCATGATTGGCTATGAAGTGGTGACCGCTGCCGACGGTGAAGAGGCGATCGCCACTTTCCATGAGAGCGACCCCGATTTGGTGGTGTTGGATGTGATGATGCCCAAGCTGGACGGCTACGGCGTTTGCCAAGAACTACGTAAAGAATCGGACATTCCCATTATTATGCTTACCGCCCTGGGGGATGTGGCCGATCGCATTACCGGTTTGGAACTGGGGGCCGATGATTATGTGGTGAAGCCCTTTTCCCCCAAGGAATTGGAAGCTAGAATCCGGTCCGTGCTACGGCGGGTGGACAAAAACGGTATGCCgcgcgctt
