## Supplementary Figures for "Optimization of *arsB* (Acr3) as a positive selection marker in the cyanobacterium *Synechocystis* sp. PCC 6803"

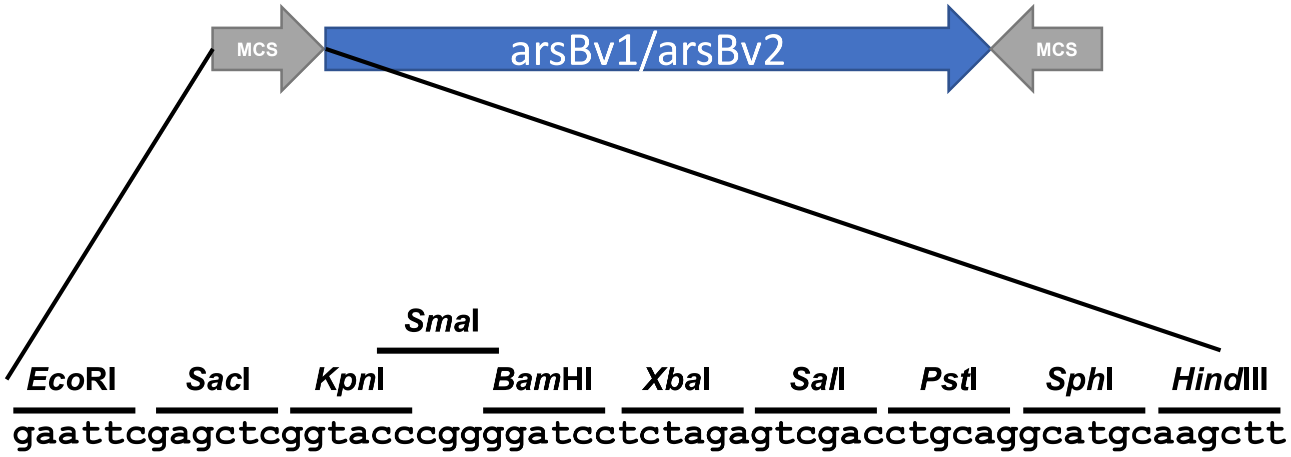


**Figure S1. pRLARSB plasmids containing the *arsB* cassettes.**
Schematic representation of the insert in pRL139 containing the *arsB* cassettes. The symmetric polylinker is also shown.

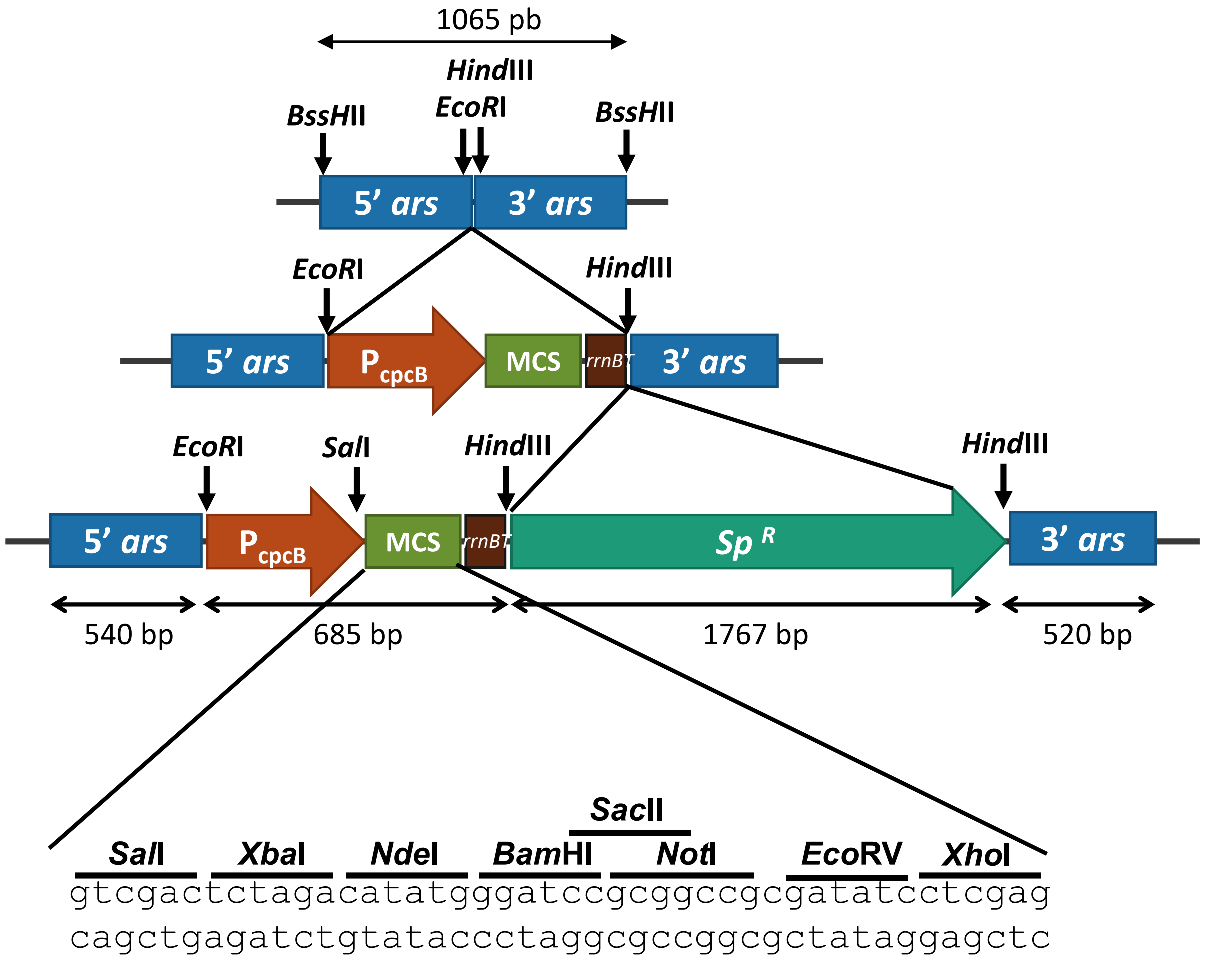


**Figure SX. Plasmid for deletion of *arsBHC* and expression of genes in Synechocystis.**
Schematic representation of the construction of pΔBHC_Sp. The plasmid includes the $P_{cpcB}$, a MCS (which sequence is also shown), *rrnB*$T_1$ and spectinomycin/streptomycin cassette and flanking region targeting the *arsBHC* locus.
